# A nanotube-mediated path to protocell formation

**DOI:** 10.1101/388405

**Authors:** Elif Senem Köksal, Susanne Liese, Ilayda Kantarci, Ragni Olsson, Andreas Carlson, Irep Gözen

## Abstract

Cellular compartments are membrane-enclosed, spatially distinct microenvironments which confine and protect biochemical reactions in the biological cell. On the early Earth, the autonomous formation of compartments is thought to have led to the encapsulation of nucleotides, thereby satisfying a starting condition for the emergence of life. Recently, surfaces have come into focus as potential platforms for the self-assembly of prebiotic compartments, as significantly enhanced vesicle formation was reported in the presence of solid interfaces. The detailed mechanism of such formation at the mesoscale is still under discussion. We report here on the spontaneous transformation of solid surface-adhered lipid deposits to unilamellar membrane compartments through a straightforward sequence of topological changes, proceeding via a network of interconnected lipid nanotubes. We show that this transformation is entirely driven by surface-free energy minimization and does not require hydrolysis of organic molecules, or external stimuli such as electrical currents or mechanical agitation. The vesicular structures take up and encapsulate their external environment during formation, and can subsequently separate and migrate upon exposure to hydrodynamic flow. This may link, for the first time, the self-directed transition from weakly organized bioamphiphile assemblies on solid surfaces to protocells with secluded internal contents.

**Significance:** The nature of the physical and chemical mechanisms behind the formation, growth and division of the earliest protocells is among the key questions concerning the origin of life. Establishing a simple pathway for the assembly of protocell structures from the primordial soup is a particular challenge. Emerging evidence supporting the assumption that solid surfaces have a governing role in protocell formation has recently expanded the scope, and created new inspiration for investigation. By presenting a physical path from self-assembled amphiphile-based membranes on solid surfaces to spherical single-membrane compartments via a consistent sequence of transformations, solely driven by the materials properties of the interfaces, a direct link between the presence of functional biomolecules and the development of protocells can be established.

## Introduction

Topologically closed membranes are among the most conserved components of living systems. The earliest cell-like constructs were likely single amphiphile-membrane enveloped aqueous microcontainers enclosing nucleotides(1, 2). In contemporary eukaryotic cells, multiple membrane compartments provide functionally specialized spaces that allow for localization and simultaneous operation of vital physiological processes that use common precursors. It has been argued that, with the ability of encapsulating nucleotides, maintaining them in physical proximity, and empowering their replication by shielding them from random mixing, such compartmentalization may have facilitated the first replication reactions, thus was a critical enabling factor for the emergence of life(2, 3).

How compartments may have developed on the early Earth, and what their exact structural and dynamic characteristics were, is currently subject of intense debate(3-5). The first compartmentalization may have occurred with the self-assembly of previously disordered constitutive molecules as a result of specific, local interactions, without any distinct external stimuli. Among a variety of eligible molecular species, amphiphiles such as fatty acids or phospholipids are strong candidates, since they can self-organize to vesicular constructs. Because of their high critical micelle concentration (CMC) and high exchange rates, fatty acids are proposed more frequently as precursors for protocell formation as compared to phospholipids. However, the synthesis of some ubiquitous components of the biological membranes such as phosphatidic acid (PA), phosphatidyl etanolamine (PE) and phosphatidyl choline (PC), has been experimentally shown to also occur under primordial Earth conditions(6-9). Primitive spherical compartments formed by self-assembled membranes share a number of characteristic features with biological cells, whose plasma membrane is also primarily composed of phospholipids. Among them are structural flexibility, wide size distribution, selective and tunable permeability as well as the ability for growth and division, and uptake and encapsulation of chemical reactants.

Most of the existing hypotheses on the formation of original biocompartments focus on bulk aqueous environments(3, 10), where a few are concentrating on assembly on solid surfaces, such as pyrites or clays(3, 4, 11, 12). The contact with surfaces would help reduce the energy barrier required for the assembly and enable the catalysis of chemical reactions. Under prebiotic conditions, organic molecules are thought to have self-assembled on inorganic solid surfaces, forming soft proto-biofilms(4), although it is argued that the strong association constants which make the membrane adsorption on solid interfaces favorable would then prevent the detachment of the assembly(3). In fact, Szostak and co-workers have made an important step ahead when they showed enhanced vesicle formation upon adsorption of sheets of amphiphiles on various types of surfaces including the minerals(3, 11, 12). The exact details of this formation process at the mesoscale remain to be elucidated.

Here we report on the spontaneous, step-wise transformation of surface-adsorbed lipid deposits into vesicular compartments. The deposits, which are giant multilamellar lipid reservoirs, undergo topological changes, sequentially adopting the forms of 2D molecular lipid films, then toroidal lipid nanotubes, and finally consistently unilamellar giant compartments. The transformation is driven by surface-free energy minimization, and does not require energy-providing chemical transformations, e.g. the hydrolysis of organic molecules, or external physical stimuli such as electrical currents or mechanical agitation. We show that the compartments encapsulate and hold a part of the medium they are created in, and can separate from the surface and migrate under the influence of hydrodynamic forces.

The observed phenomena provide strong evidence that the transition from molecular amphiphile films on solid surfaces to spherical protocell compartments can occur spontaneously in simple aqueous environments in a materials properties-driven, rather than a chemically initiated, process.

## Results

### Spontaneous emergence of giant vesicular compartments from surface-adhered lipid nanotubes

We start the experiments by placing lipid reservoirs (multilamellar vesicles, MLVs) onto a SiO_2_ substrate, which results in self-spreading of each reservoir on the solid support as a circular double bilayer membrane(13, 14). Minutes after the MLV deposition, due to the continuous adhesion of the circular membrane’s periphery to the solid substrate, the membrane tension reaches lysis tension (5-10 mN/m), leading to rupturing(14). On the ruptured lipid patches, we observe the formation of numerous (100-1000 per patch) giant vesicular compartments, originating from surface-adhered phospholipid nanotubes (Fig. 1a-b). The tubes, the formation of which is described in detail in subsequent paragraphs, appear during the retraction of the distal (upper) bilayer off the proximal (lower) bilayer of a double bilayer membrane. The retracting upper membrane leaves behind a network of randomly pinned lipid nanoconduits on the proximal bilayer (Fig. 1a). Islands of remaining distal membrane can be observed in Fig. 1b. The vesicles emerge from the nanotubes, initially adopting a dome-shape(Fig. 1c-e), and over time, a spherical giant unilamellar vesicle (GUV) (Fig. 1f-h). The buds (d≥0.8µm, Mov. S1-2) appear within minutes, and mature over a few hours to larger vesicles with diameters of 5-10 µm. During vesicle growth, the nanotubes are connected to the vesicles on one end, and on the other end to the proximal bilayer (Fig.1e).

**Figure 1.**
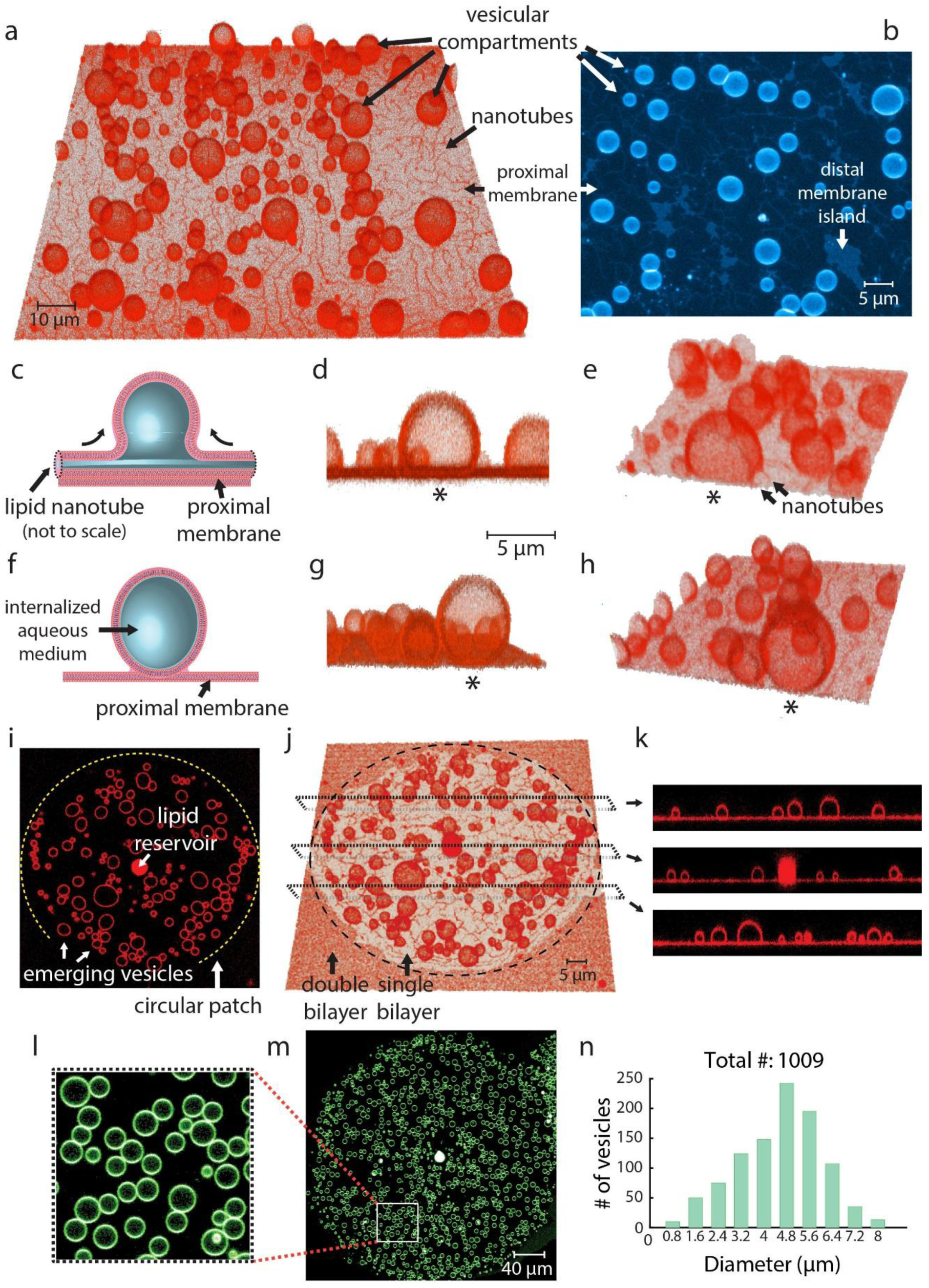
Giant vesicular compartment formation from surface-adhered lipid nanotubes. **(a-b)** Confocal micrographs, reconstructed in 3D, showing supported vesicles from **(a)** tilted view and **(b)** top view. Vesicles emerge from lipid nanotubes, which spontaneously form during de-wetting of the proximal (lower) bilayer by the distal (upper) bilayer of an initially intact double bilayer membrane. **(c-e)** semi-grown and **(f-h)** fully-grown vesicular compartments. The vesicles, most of which are pinned to the proximal membrane via nanotubes adopt a dome-shape **(c-e)**. The connection of the vesicle to nanotubes, is captured in **(e)**. Over time, compartments mature to a spherical form **(f-h)**. Vesicles which are identical in **(d-e)** and **(g-h)**, respectively, are marked with an asterisk. **(i-k)** Confocal micrographs of a circular membrane patch accommodating several vesicles: **(i)** top view (x-y plane), **(j)** tilted 3D view (x-y-z plane), **(k)** side views (x-z plane). The three separate profiles depicted in **(k)** correspond to the regions in **(j)** marked with dashed lines. The circular patch, as described in the main text in detail, is a result of self-spreading of a multilamellar lipid reservoir. **(l-m)** Fluorescence micrograph of vesicles emerging from a circular patch, imaged in the x-y plane. In **(l)**, the region in **(m)** framed with a white line, is magnified. **(n)** Graph showing, the number of vesicles in the membrane patch in **(m)**, versus vesicle diameter. The patch contains over a thousand vesicles in total.

When an initially circular double membrane ruptures and transforms into a single bilayer, the circular contour is largely maintained. The vesicles therefore are concentrated in circular regions, with MLVs in the center (Fig. 1i-k). Such vesicles, emerging from a ruptured circular patch, are shown in Fig. 1i and j from different angles. In Fig. 1k, three sections from panel j (frames in dashed lines), have been shown from the profile view. Except for the MLV, the vesicles appear to be unilamellar. If MLVs are added in high numbers to increase the membrane coverage of the surface, occasionally, the spreading double bilayer membrane of adjacent reservoirs can spread around a ruptured single bilayer patch, as shown in Fig. 1j.

A large lipid patch (d=∼300 µm) can accommodate about a thousand unilamellar vesicles (Fig. 1 l-n). The vesicle sizes vary from 1 to 10 μm in diameter, most in the range of 4-6 µm.

### Formation of nanotubes and vesicles

During the rupturing described above, the distal membrane continuously de-wets the proximal, leaving behind nanotube threads (Fig. 2a-c). While the edge of the bilayer is shown open in the schematic drawings, in reality it is closed (inset Fig. 2a), resulting in an increasing edge energy cost(15, 16). The de-wetting process at a peripheral region of a lipid patch, leading to the formation of nanotubes and vesicles, is depicted in Fig. 2b-c, Mov. S3. (*cf*. Mov. S4-5 for additional movies of growth). The contour of the edge of the retracting double bilayer membrane in Fig. 2b, is superimposed on Fig. 2c (yellow dashed line). Pinning among bilayers, or between the bilayer and the solid substrate is mediated by the Ca^2+^ in the ambient buffer(13, 14, 17).The elongation of the edges of the pinned threads due to continuous retraction of the distal bilayer causes the increase of edge tension, which eventually must lead to bending and toroidal tube formation (Fig. 2d-f). Small buds appear on these tubes which form and reside on the proximal membrane (Fig. 2 g-h). The buds initially adopt a semi-spherical, or dome-like shape with elongated sides which are connecting to the nanotubes. These semi-spherical buds develop into spheres over time. It can be seen that the regions which are saturated with vesicles have a reduced nanotube density (Fig. 2i-j), indicating that the tubes are consumed during vesiculation.

**Figure 2.**
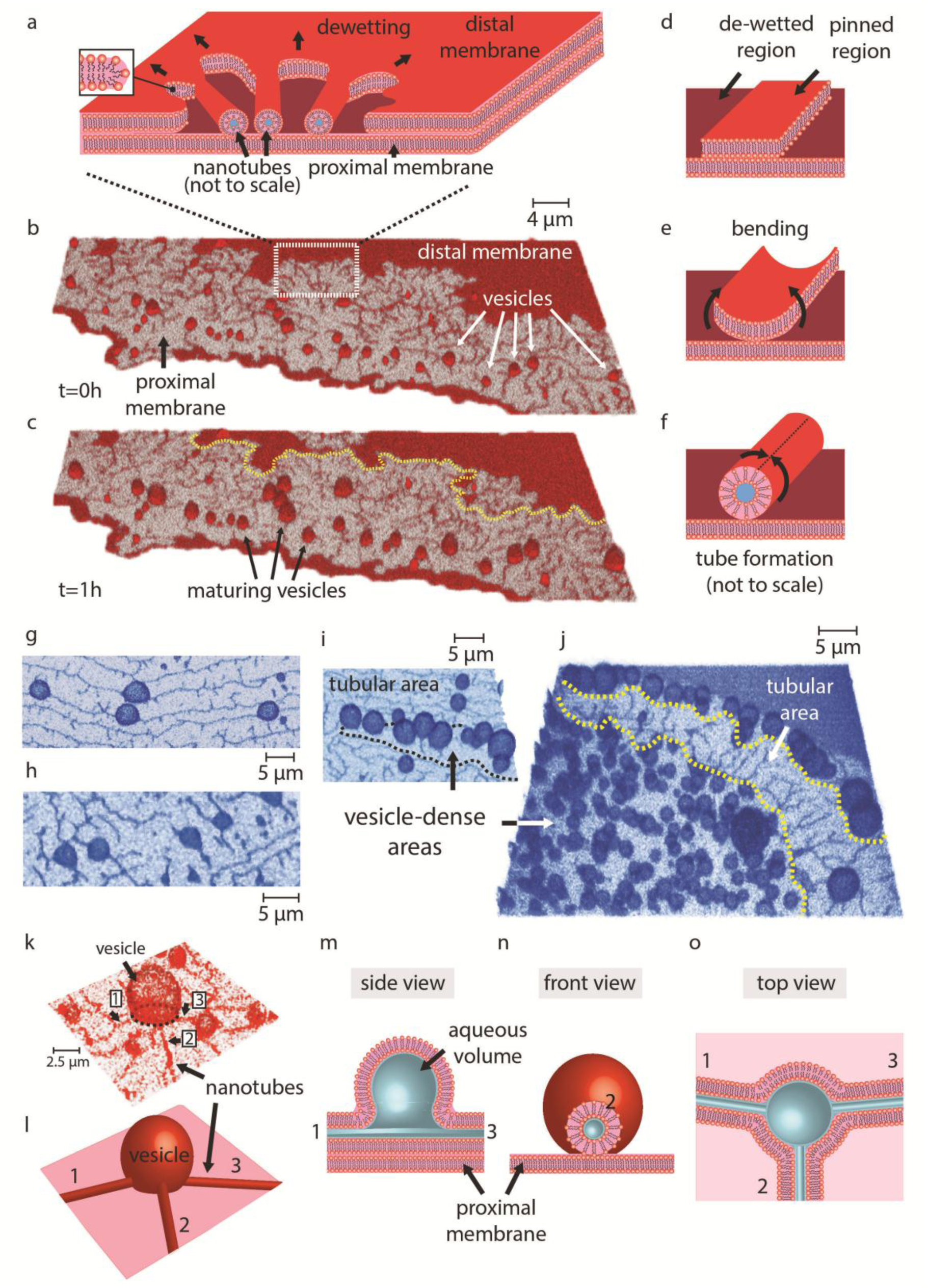
Formation of nanotubes and vesicles. **(a)** Spontaneous formation of lipid nanotubes during de-wetting. Schematic representation of retraction of the distal membrane (red) over the proximal membrane (burgundy), leaving behind lipid threads. The confocal micrographs showing the corresponding experimental process, are depicted in **(b-c)**. The yellow, dashed line in **(c)** represents the contour of the distal membrane in **(b)**. Initially, the parts of the distal membrane which are pinned to the proximal one, remain **(d)**. Such thin threads growing in length as the de-wetting continues, create increasing edge tension which favors bending **(e)** followed by the nanotube formation (**f**). The nanotubes are not drawn to scale as the diameter of a lipid nanotube is 100-200 nm. In the cartoons, the membrane edge is shown as open. In reality, the membrane edge is curved (inset to **(a)**), to avoid the exposure of hydrophobic moieties of the membrane lipids to the aqueous buffer. The vesicles emerging from the nanotubes in **(b)** have grown larger in **(c)**. **(g-h)** Emerging vesicular buds on lipid nanotubes. The vesicles appear spherical in the middle and elongated on the sides, where they connect to the nanotubes. **(i-j)** The areas populated by vesicles (regions framed in dashed lines) contain relatively less or no tubes. **(k)** The confocal image, reconstructed in 3D, and **(l)** the corresponding schematic representation, of a vesicle emerging from/connected to, multiple lipid nanotubes (numbered). **(m-o)** Schematic drawings explaining the configuration of the vesicle and nanotubes from different views.

The vesicular compartments occasionally appear at the junction of several nanotubes. The confocal micrograph of a vesicle connected to at least 3 tubes (numbered), is shown in Fig. 2k, and the corresponding schematic representations in Fig. 2l-o. Fig. 2m shows the profile view along the nanotube #1-the vesicle-the nanotube #3. In Fig. 2n, nanotube #2 is elongating perpendicular towards the plane of the page. The vesicle appears behind the cross-sectional view of the tube. Fig. 2o is a top view of the vesicle and the three tubes with the proximal bilayer underneath (light pink color).

### Consistent unilamellarity

Because of the formation mechanism, the vesicles originating from a single bilayer, appear consistently unilamellar. A confocal micrograph of a membrane region capturing several small vesicles and nanotubes on a proximal bilayer, is shown in Fig. 3a. The plots in Fig. 3b-d show the fluorescence intensity profiles along the orange arrows in Fig. 3a in indicated directions (for additional number of analysis *cf*. Fig. S1). The focal depth (x-z) of the confocal image shown in x-y plane is ∼1 µm. This means, the micrograph of a vesicle on the surface with a diameter of ∼2 µm, captures the fluorescence signal starting from the solid substrate until only 1 µm above, which corresponds to approximately half of the spherical vesicle. For a 5 nm thick proximal bilayer and the lipid nanotubes (d=100-200 nm), the fluorescence signal of the entire structure can be collected in x-z direction. Accordingly, the intensities across the proximal membrane (single bilayer), the equator of the vesicles (two bilayers) and the nanotubes (three bilayers) appear in a 1:2:3 ratio (Fig. 3b-d). To complement the analysis in Fig. 3a-d, and to verify the consistent lamellarity of multiple vesicles, a false colored confocal image of a membrane area containing several vesicles is shown in Fig. 3e-h. The z-axis of the scan is divided into intensity levels of an 8bit image (255 levels). The emission from labeled membranes of more than a hundred vesicles has nearly identical intensity (single bilayer) represented by the green color (Fig. 3e). The regions between adjacent vesicles with overlapping bilayers appear as blue (*cf*. false color intensity scale), corresponding to the double intensity of a double bilayer structure (Fig. 3f-h).

**Figure 3.**
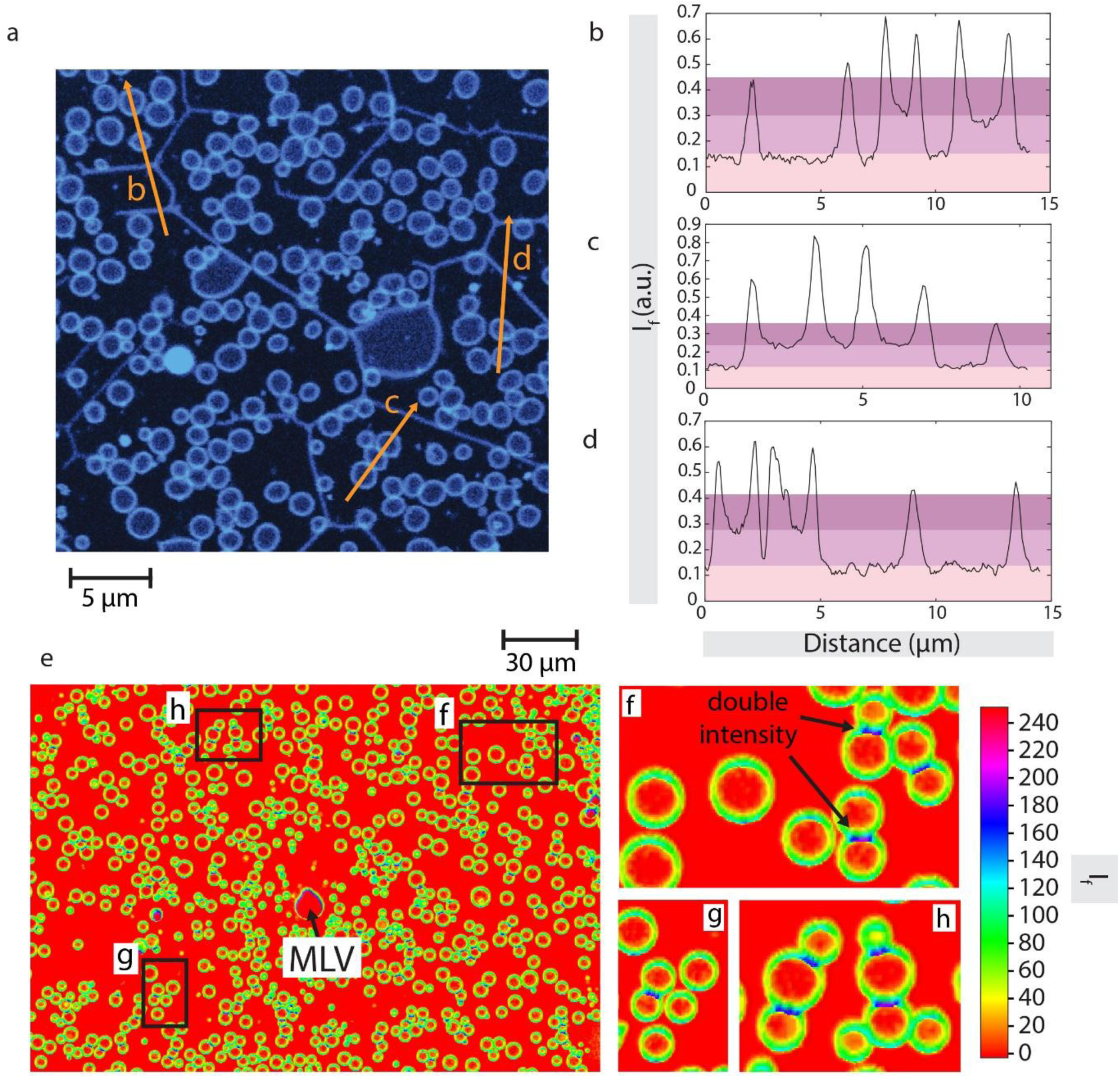
Lamellarity analysis. **(a)** Confocal fluorescence micrograph of a membrane region in x-y plane. **(b-d)** The fluorescence intensity profiles along the orange lines in **(a)**. The arrows indicate from left to right, the direction of the profiles in panels **(b-d)**. The intensities across the proximal membrane (single bilayer), the equator of the vesicle (two bilayers) and the nanotube (three bilayers) appear in a 1:2:3 ratio. **(e)** The false-colored fluorescence intensity plot of a membrane region containing over 100 vesicles shows nearly identical intensity (single lamellarity). **(f-h)** Close-ups of 3 different regions from **(e)**. The overlapping membranes, where adjacent vesicles come in contact, have twice the intensity of the non-overlapping regions, indicating single and double bilayer structures.

### Numerical simulations of vesicle nucleation

In order to determine the relationships between the membrane area, its membrane bending energy, and the distance between two pinning points, we carried out finite element simulations (Fig. 4), using the software *Surface Evolver* (SI). The pinning sites, or membrane defects, are assumed to be immobile points along the tube (Fig. 4a) (*cf.* discussion). The ratio of the distance between the two closest pinning points *l*, and the tube radius *r* gives the dimensionless number 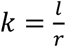, which is varied systematically in the simulations. In each simulation for a given *k*, a fixed membrane area *A* > *A*_*cylinder*_ was defined, and the membrane area was allowed to equilibrate to a shape that corresponds to a minimum in bending energy *E*. In order to normalize the results for different distances between pinning defects, *E* and *A* were divided by their respective values for a cylindrical tube, with *E*_*cylinder*_ = 2*πk* and *A*_*cylinder*_ = 2*πrl*. Fig. 4a shows snapshots from the simulation of a nanotube evolving to a vesicle for *k* = 20. The model predicts the formation and growth of vesicles similar in size and shape to the ones observed experimentally (Fig. 2g-h). Fig. 4b shows the relation between the total energy and the surface area of the adhered tubes for different values of *k*. For *k* ≤ 15 the energy exhibits a minimum, i.e., the tube would have to overcome an energy barrier (energy of the highly curved neck region at either end of a bud) in order to transform into a spherical vesicle. In contrast, for longer tubes (*k* ≥ 20) the energy decreases monotonically, which facilitates vesicle growth. Vesicle budding is initiated at 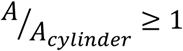 where the total energy starts to deviate from the energy of a free-floating tube (solid black line). For large surface areas 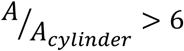, the energy approaches the value of a spherical vesicle (*cf.* SI). For each *k* value, the dashed lines in Fig. 4b point to the magnitude of the total bending energy of spherical vesicles that are free of adhesion.

**Figure 4.**
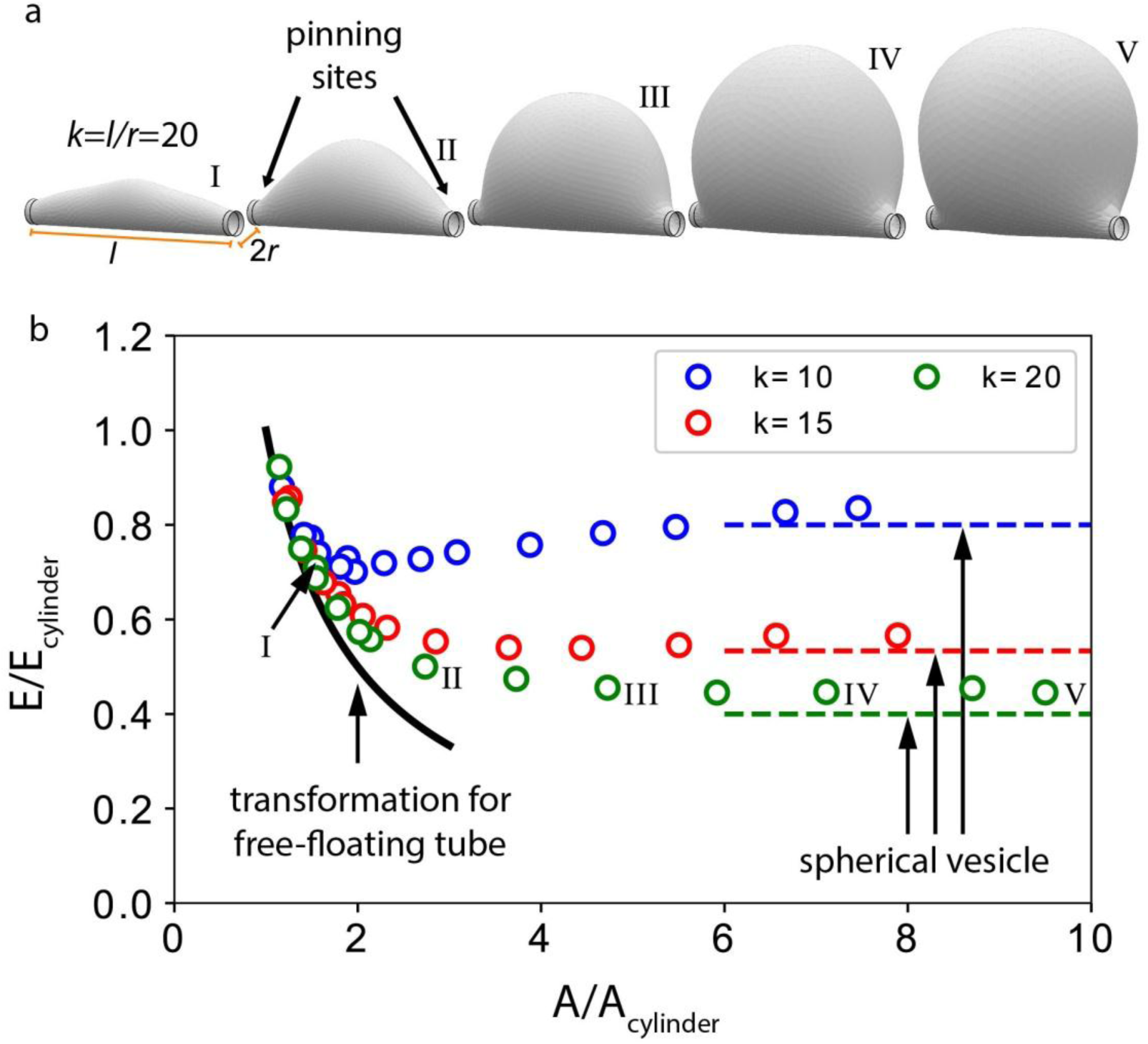
Bending energy E during tube-vesicle transformation. **(a)** Simulation snapshots showing the evolution of a nanotube to a vesicle, where the ratio *k* of the distance between two immobilized points along the tube *l* to tube radius *r*, is 20. **(b)** Plots showing bending energy vs. surface area of the adhered membrane structure (tube, bud, or vesicle) for three different values of *k*. For *k* < 20, indicating a short distance between the two fixed points (pinning sites), tube-to-vesicle transformation is unfavorable, while for *k*≥ 20, vesicle formation is favored. The energy values labeled I-V correspond to the vesicle shapes shown in **(a)**. The corresponding dashed lines indicate the surface energy of completely spherical vesicles that are free of adhesion. The black solid line represents the transformation of a free-floating tube to a vesicle. Each simulation is repeated at least nine times for different initial conditions and the variation in energy is within the size of the markers.

### Encapsulation

Next, the supported bilayers were transformed in HEPES buffer containing fluorescein. Subsequently, the external solution was exchanged with HEPES buffer of identical composition and pH, but this time free of fluorescein (Fig. 5a). The resulting vesicles encapsulated fluorescein in their internal volume, and maintained it within. The cross section from the profile (x-z plane) of several vesicles is shown in Fig. 5b. Fig. 5c-r show confocal micrographs of 4 membrane regions from different lipid patches populated with vesicles encapsulating fluorescein and the corresponding intensity profiles. The vesicle membranes are labeled with rhodamine dye, (red, Fig. 5c, g, k, o) and the vesicles encapsulate 0.1 mM fluorescein (green, Fig. 5d, h, l, p). Fig. 5e, i, m, q show the rhodamine and the fluorescein channels superimposed. The fluorescence intensity profiles along the arrows in Fig. 5e, i, m, q are shown in Fig. 5f, j, n, r; respectively. In these plots, the vesicular membrane (plot in red color) appears as two close-spikes, each corresponding to the bilayer membrane of the vesicle; and the fluorescein signal (plot in green color) remains in between the spikes, corresponding to the interior of the vesicle.

**Figure 5.**
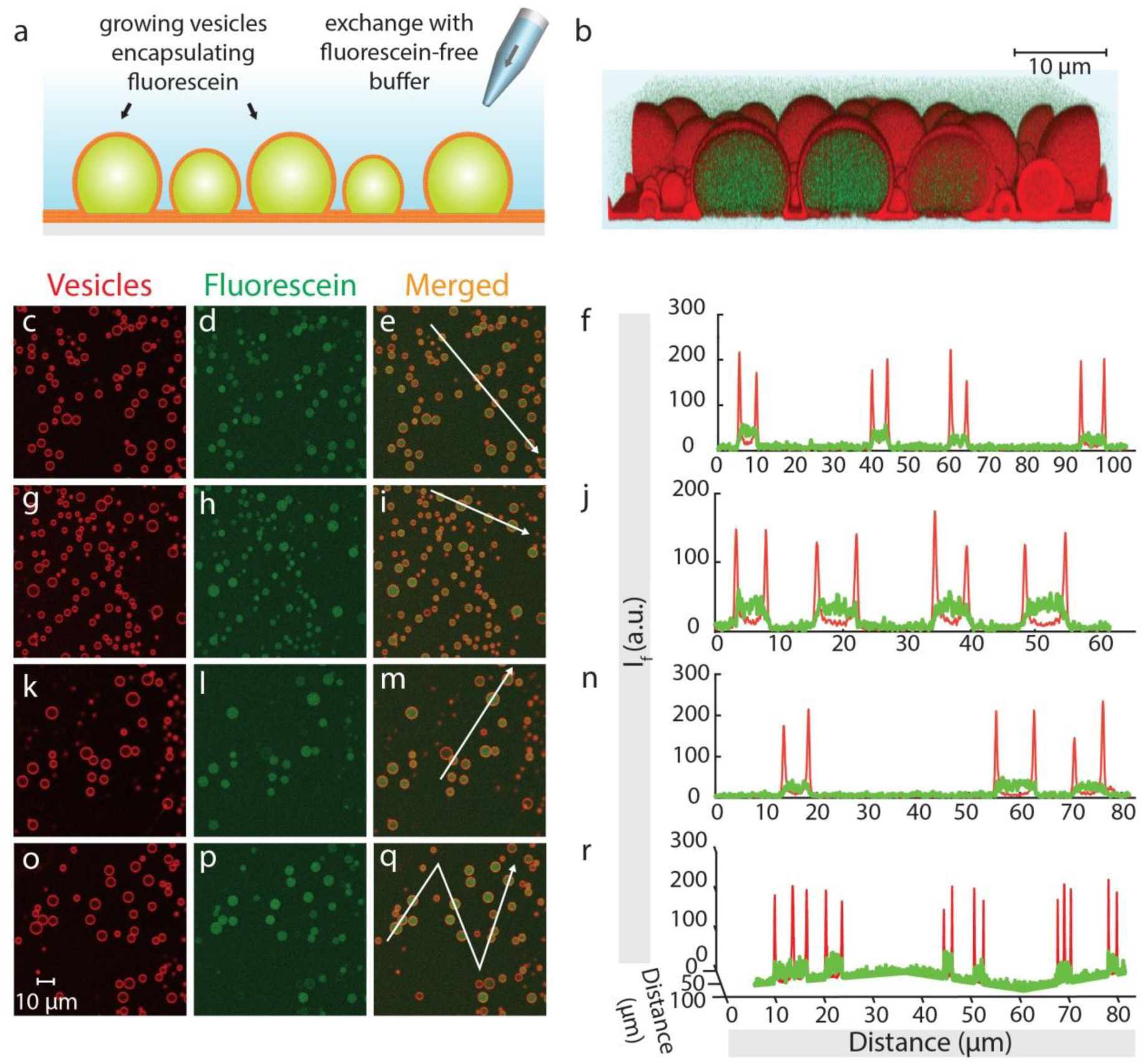
Encapsulation of fluorescein in vesicular compartments. **(a)** Schematic illustration and confocal micrograph of vesicles encapsulating fluorescein (Fl), from side view (x-z plane). Vesicles were grown in Fl -containing HEPES buffer. After maturation, the ambient solution was exchanged with Fl -free HEPES buffer, using an automatic pipette. **(c-r)** Fluorescence micrographs (x-y plane) and corresponding intensity profiles of vesicles encapsulating Fl. Panels **c-g-k-o** show membrane fluorescence, **d-h-l-p** the encapsulated Fl; and **e-i-m-q** combine both. The graphs in **f-j-n-r** show fluorescence intensity along the white arrows in **e-i-m-q**. The arrows indicate the direction of the depicted profiles from left to right. Red color represents the membrane fluorescence, and green color fluorescein.

### Separation and migration of vesicles

The GUVs which were initially anchored to the supported bilayer were exposed to hydrodynamic flow in the range of 10-100 nl/s by using the ‘multifunctional pipette’, an open-space microfluidic device(18, 19). This exposure led to separation of the vesicles from the assembly (Fig. 6). Briefly, the pipette injects a fluid stream into the vicinity of the vesicles through a central channel, while two peripheral channels simultaneously aspirate, creating a fluid recirculation zone within the existing medium (ambient buffer) without mixing the two (Fig. 6a). The effective hydrodynamic flow in the exposure zone causes loosely surface-adhered objects to separate, and they can be collected in on-chip wells. By means of this setup, the surface-adhered vesicles were separated from the membrane patch, collected and subsequently transferred with an automatic pipette onto gold-coated glass substrates in order to avoid rapid adhesion and rupturing, which typically occurs on glass slides. The vesicles adhering to the gold substrates are shown in Fig. 6b and 6c (x-y-z plane). The inset to Fig. 6c shows the top view (x-y) of vesicles #1-4. Fig. 6d-g and 6h-j show two individual experiments capturing the separation and migration process. To be able to visualize the recirculation zone, fluorescein was used as tracer prior to the migration experiments (Fig. 6d, dashed lines).

**Figure 6.**
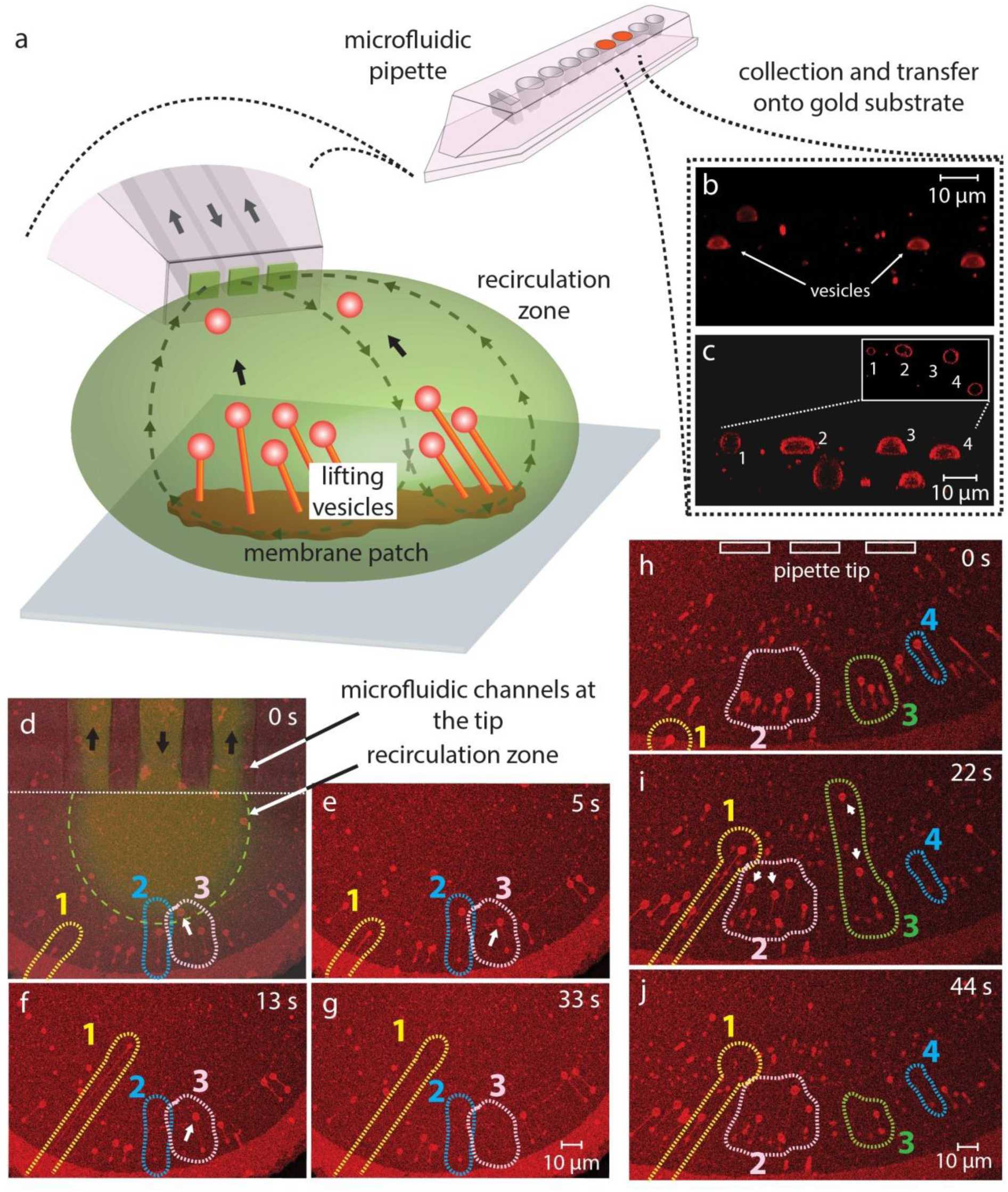
Separation and migration of vesicles after exposure to hydrodynamic forces. **(a)** Schematic illustration of the collection process, utilizing an open-space microfluidic device. The device circulates a liquid at its tip (green), by injecting from a central channel and aspirating simultaneously from the two peripheral channels. The objects in the ambient buffer entering into the recirculation zone separate from the membrane and migrate to collection wells, from which they can be retrieved. **(b-c)** Confocal florescence micrographs of the collected vesicles placed on gold substrates(x-y-z). Inset to **(c)** shows four of the vesicles in **(c)** (numbered) in x-y plane. Fluorescein as tracer was used in **(d)** for better visualization of the recirculation zone (dashed line). **(d-g & h-j)** show separation of initially membrane-adhered vesicles from two different lipid patches. Transient nanotubes form while pulling the vesicles into the pipette. The vesicles in regions 1-3 in **(d-g)** and in region 1 and 4 in **(h-j)** were collected. A few vesicles in regions 2-3 in **(h-j)** remain on the membrane after recirculation is terminated.

Our initial attempts of exposing the vesicles to several cycles of aspiration with the microfluidic pipette were resisted by the strong anchoring of the vesicles to the bilayer underneath (not shown). To be able to facilitate separation and collection, we weakened the pinning/anchor points of the vesicles. Accordingly, we exchanged the Ca^2+^-containing ambient buffer with a Ca^2+^ chelator-containing solution. (1,2-bis (o-aminophenoxy) ethane-N,N,N′,N′-tetraacetic acid (BAPTA) and Ethylenediaminetetraacetic acid (EDTA)(18, 20) were used in the experiments, and the vesicles with reduced pinning due to Ca^2+^ depletion were easily pulled into the pipette. During collection, some of the vesicles which were lifted by the aspiration force, remain connected to the lipid patch through a nanotube (Fig. 6d-g/ Mov. S6; 6h-j/ Mov. S7). Extrusion of nanotubes by exposing the vesicles to hydrodynamic flow is a well-characterized phenomenon(21). All vesicles in regions 1-3 in Fig. 6d-g and in regions 1 and 4 in Fig. 6h-j separated and migrated towards the collection wells, while a few vesicles in regions 2-3 in Fig. 6h-j remained on the membrane after recirculation was terminated.

## Discussion

### Tube formation

During double bilayer membrane rupturing, the distal membrane rapidly and continuously de-wets the proximal membrane. Throughout this process, some regions of distal membrane remain on the proximal membrane, owing to pinning. The distal membrane regions left-behind during de-wetting commonly appear in the form of tubular threads. A thread forms between the retracting membrane and a pinning site. Very rapidly, the edge energy *E*_*edge*_ = *γl*, where *γ* is the line tension (5-10pN) and *l* is the length of the elongating ruptured edge(15, 16), increases until the edges of the thread eventually bend, leading to toroidal tube formation. Fig. 2d-f show one possible mechanism for nanotube formation, which assumes that the pinning creates a flat interface between the proximal and the distal bilayers. The lateral peripheries of a growing distal membrane thread rapidly bend upwards and form a closed structure.

Pinning sites can form due to adhesion of two bilayers in close proximity caused by Ca^2+^ ions(13, 14, 17, 22-24). The pinning sites can also form dynamically due to rapid alterations in membrane tension(25-27). The membrane patches we observe are large, with diameters of 300-400 µm. In such large areas, the membrane relaxation, for example due to a pore opening, cannot instantly propagate through the membrane evenly, but rather influences the membrane tension locally. Local membrane tension may indeed vary throughout our experiments, indicated by the simultaneous collapse and emergence of vesicles on the same membrane region (Fig. S7). In our system the membrane locally experiences regions of strong and weak adhesion due to pinning and due to fluctuating membrane tension. The interaction of a membrane nanotube with a bilayer has been previously studied in great detail with dissipative particle dynamics (DPD) simulations(25), which predict that strong adhesion leads to merging of the tube with the underlying bilayer membrane, whereas weak membrane adhesion promotes pearling(25). Pearling instabilities can be observed when the local tension in the tube compared to the membrane material around is high(28, 29). When excess membrane material becomes available in the membrane retraction process, the tube, in order to reduce its surface energy, transforms into a string of small spheres which are connected with very thin membrane necks.

### Vesicle formation and growth mechanism

The experimental evidence shows that the vesicles are formed via enlargement of the lipid nanotube fragments residing on a proximal bilayer (Fig. 1 and Fig. 2). Contraction of lengthy (*l*=100 µm) lipid nanotubes to giant vesicles in short time scales (seconds to minutes) requires significant flip-flop rates, as well as rapid influx of ambient buffer. While high flip-flop rates can be accommodated during rapid shape transformations in high curvature systems such as nanotubes(30), the fast influx of an aqueous solution to vesicle is only possible through membranes with abnormally high permeability(31). Contraction of free-floating nanotubes to vesicles in short time scales therefore, result in stomatocyte-like vesicular structures containing folded double bilayers(31). In our experiments, the nanotubes are pinned to the surface underneath. The pinning prevents rapid contractions which would lead to stomatocyte-like vesicles mentioned above. The formation of small buds in the micrometer range occurs within minutes, during which inter-leaflet lipid transfer might occur, whereas the maturation to giant vesicles takes hours to days, pointing to a simple mechanism of tube inflation in a low membrane tension regime(32). Long time durations enable lateral lipid migration between the nanotube and the vesicular membrane. The lamellarity analyses shown in Fig. 3 confirm that all of the resulting vesicle membranes are single bilayers, which indicates that the vesicles draw lipids from a remote single-bilayer source via a membranous connection, in contrast to the typical osmotically driven swelling processes of multilamellar reservoirs.

### Source of material for vesicle formation

The material consumed in the vesicle formation process can be efficiently supplied by the tubes. To form a vesicle with d_vesicle_=5 µm, a nanotube (d_tube_=100 nm) with a length of 250 µm is required. This is well within the range of the tube lengths we are observing(20). To fill the interior volume of this vesicle, however, solely from the internal aqueous volume of a toroidal lipid nanotube, the required tube length would be ∼8 mm per vesicle. A significant portion of the aqueous volume inside the vesicles is therefore likely supplied from the ambient buffer through defects or transient pores in the vesicle membrane. Inflation of a toroidal nanotube by means of microinjection of an aqueous medium through a membrane opening, resulting in a giant lipid container, has been reported earlier(32, 33). In these studies, the influx of 50×10^−15^ l/s of injected aqueous medium controlled by the micropipette was reported, corresponding to 10-60 µm^2^/s of membrane replacement(32). Our observations, showing the spontaneous inflation of the tube to a 5 µm vesicle, indicate the influx to be as small as 2×10^−18^ l/s, which corresponds to a membrane replacement of 2×10^−3^ µm^2^/s. This rate is well-below the values observed for the micropipette injection method, supporting that the tube swelling/inflation as a mechanism for vesicle formation is realistic.

We observe that the entire vesicle population internalizes fluorescein (Fig. 5). This further indicates that the internalization process involves primarily membrane openings, through which the external fluid can enter. This is an important detail, since the encapsulation of fluid could in principle also proceed through osmotic pressure-induced swelling, where water molecules penetrate a bilayer membrane in order to relax an ion concentration gradient. In this case, larger molecules would not be internalized along with the water. Since the compartments maintain integrity and hold the internalized contents, the pores must close after vesicle formation. It is likely that the closure of the transient openings terminate vesicle growth.

### Driving force for tube-to-vesicle transformation

Lipid membrane nanotubes are highly curved. From the energy perspective, they are disfavored compared to spherical vesicles. By transforming a nanotube to a spherical vesicle, the curvature of the membrane is reduced and the surface free energy of the membrane is minimized(34). Note that the surface free energy of a membrane is not exclusively defined by the bending energy, but it is the dominating term when high curvature is involved. When membrane material becomes available due to rupturing, the highly curved tube relaxes by incorporating the excess material, spontaneously transforming into the less curved, energetically more favorable spherical shape. The experimental observations (Fig. 2g-h) and the predictions by the model (Fig. 4) are in agreement. Vesicle formation is to the most extend driven by membrane curvature and bending energy minimization; both are materials properties-related features and processes.

### Location of nucleation of the vesicles

We currently do not know the factors which determine the locations of the nucleation sites. It is conceivable that the pinning sites, which are essential for tube formation, can also influence vesicle budding. One possible mechanism governing vesicle formation involves that pinning sites or membrane defects are also nucleation sites. Some vesicles are indeed connected by multiple tubes (Fig. 2), which suggests that at least one pinning site is situated beneath the vesicle. Pearling instabilities in between two pinning sites, arising from a tension gradient between tube and surrounding membrane material, would provide a nucleation site, from which the vesicles can evolve. If we presume that the pinning points have a role in vesicle nucleation, through the numerical simulations we obtained the minimum distance of two points in between which a tube-to-vesicle transformation can occur. Bud formation initiates at *k* = 20 (Fig. 4b), i.e., for d=100 nm the defects must be at least ∼1 µm apart from each other. This corresponds to the smallest vesicle sizes we observe (Fig. 1n). Note that our model assumes that the vesicles nucleate in between two pinning sites. These sites, however, may not be necessarily physical boundaries of vesicle growth. Since number and density of the pinning sites are not exactly known, it cannot be excluded that vesicles incorporate several defects during growth.

### Possible implications for the origins of life

We show the spontaneous formation of consistently unilamellar lipid vesicles from surface-adhered phospholipid nanotubes. These spherical compartments possess an intact lipid bilayer, and can engulf an aqueous volume containing solutes and organic compounds, both of which are essential features of a primitive protocell(1, 2). Compartmentalization is considered to be a fundamental step for the emergence of life, but how such containers were formed on the early Earth, and what their exact structural and dynamic characteristics could be, is currently subject of an intense debate(3, 5). Under prebiotic conditions, structurally and functionally distinct organic molecules, in particular amphiphiles, are thought to have self-assembled on inorganic solid surfaces, forming soft proto-biofilms. The surfaces would assist the surfactants in overcoming the energetic barrier of self-assembling in a bulk aqueous solution without support(4). Szostak and coworkers have reported facilitated vesicle formation in the presence of various types of solid particles including minerals(12), and proposed that the amphiphilic molecules were assembling into sheets close to the particle surface. The resulting vesicles were shown not to contain the fluorescently labeled layer adsorbed directly on the particles, *i.e.*, the vesicles were not physically stemming from the membrane directly in contact with the solid surface. How this enhanced formation occurred at the mesoscale, is currently not known in detail; but the proposed models(3, 12) support our findings that a second, distal bilayer forms the compartments. We also show that moderate hydrodynamic flow is able to separate and displace the vesicles from the surface, which could represent a primitive means of migratory mobility of intact protocells under reasonable ambient conditions.

We provide, for the first time, direct evidence showing a physical path of transformation from self-assembled amphiphile-based membranes on solid surfaces to spherical single-membrane compartments, proceeding via intermediate nanotubular structures. Very few assumptions have to be made to link this self-driven phenomenon directly to protocell formation. Strictly required are the presence of sufficient amphiphilic material, a high-energy solid surface and suitable ambient conditions. It has been earlier shown that upon mild agitation the amphiphilic nanotubes form multiple daughter vesicles in a more efficient way, compared to the division of a large spherical vesicle into smaller vesicles(8, 35). This observation is fundamental for the system we report here, and we present strong evidence that due to the intrinsic physical and materials properties of the system, even without the need of agitation, sonication, mechanical or hydrodynamic disturbance, nanotubes will spontaneously give rise to vesicular compartments.

## Methods

### Stock lipid suspension

Various recipes of lipid mixtures were tested, a full list of which is provided in Table S1. To prepare each lipid composition, dehydration and rehydration (gentle hydration) method described by Karlsson *et al*.(36) was followed. Briefly, lipids and lipid-conjugated fluorophores in designated ratios were mixed in chloroform reaching to a total concentration of 10 mg/ml. 300 µl of this solution was placed in a 10 ml round bottom flask and the chloroform was removed in a rotary evaporator at reduced pressure (20 kPa) over a period of 6 hours. The dry lipid film at the walls of the flask was rehydrated with 3 ml of PBS buffer containing 5 mM Trizma Base, 30 mM K_3_PO_4_, 30 mM KH_2_PO_4_, 3 mM MgSO_4_*7H_2_O and 0.5 mM Na_2_EDTA. The pH was adjusted to 7.4 with H_3_PO_4_. The rehydrated lipid cake was kept at +4 °C overnight. In the final step, lipid cake was sonicated for 10 seconds at room temperature to induce the formation of giant vesicles of varying, mainly multiple lamellarity.

### Multilamellar reservoir formation

4 µl of each lipid suspension was placed on a cover slip and dehydrated in an evacuated desiccator for 20 min. The dry lipid film was rehydrated with ∼1 ml of HEPES buffer containing 10 mM HEPES and 100 mM NaCl (pH= 7.8, adjusted with NaOH) for 10 min to allow formation of MLVs. MLVs were then transferred onto a SiO_2_ substrate containing HEPES buffer with 10 mM HEPES, 100 mM NaCl and 4 mM CaCl_2_ (pH= 7.8, adjusted with NaOH), leading to spreading, rupturing and formation of vesicles.

### Surface fabrication & characterization

SiO_2_ deposited onto the glass substrates by reactive sputtering, using a MS 150 Sputter system (FHR Anlagenbau GmbH) or E-beam and thermal PVD using EvoVac (Å ngstrom Engineering), to final film thicknesses of 10 to 100 nm. The results were indifferent to film thickness. The thickness of the films was verified by ellipsometry (SD 2000 Philips). No pre-cleaning was performed before deposition. Surfaces were stored at room temperature until use. 10 nm Au films were deposited on glass cover slips on top of a 2 nm TiO_2_, using MS 150 Sputter system (FHR Anlagenbau GmbH).

### Microscopy imaging

A confocal laser scanning microscopy system (Leica SP8, Germany), with HCX PL APO CS 40x and 60x oil objectives and 20x air objective were used for acquisition of confocal fluorescence images. The utilized excitation/emission wavelengths for the imaging of the fluorophores, were as following: ex: 560 nm, em: 583 nm for rhodamine, ex: 488 nm, em: 515 nm for fluorescein, ex: 544 nm, em: 571 nm for TopFluor^®^ TMR. The results were indifferent to fluorophores.

### Encapsulation

The main experiment the details of which was described above was reproduced in HEPES buffer containing 10 mM HEPES, 100 mM NaCl and 4 mM CaCl_2_, 100 µM Fluorescein (NaFl salt, Sigma Aldrich). After vesicle formation, the excess dye in the chamber was replaced by gently exchanging the Fl-containing HEPES buffer with the Fl-free HEPES using an automatic pipette.

### Separation and migration of GUVs

An open-volume microfluidic device/pipette (Fluicell AB, Sweden) was used to expose the initially surface-adhered vesicles to hydrodynamic flow, inducing their separation and migration. The pipette was positioned by a 3-axis water hydraulic micromanipulator (Narishige, Japan). 10 mM HEPES buffer containing 100 mM NaCl, 10 mM EDTA and 7 mM BAPTA (pH=7.8, adjusted with NaOH) was used to weaken the Ca^2+^ mediated adhesion of the vesicles to the underlying membrane. The vesicles were retrieved from the waste wells of the microfluidic device using an automatic pipette and placed on Au-coated glass surfaces.

### Image processing/analysis

3D fluorescence micrographs were reconstructed using Leica Application Suite × (LasX) Software (Leica Microsystems, Germany). Image enhancements to fluorescence micrographs were done with NIH Image-J Software and Adobe Photoshop CS4 (Adobe Systems, USA). Schematic drawings and image overlays were created with Adobe Illustrator CS4 (Adobe Systems, USA). Vesicle counting and size distribution analyses were done using NIH Image-J Software. Fluorescence intensity profiles were drawn in Matlab R2017a after applying median filtering. The false-colored fluorescence intensity plot was prepared with NIH Image-J Software Interactive 3D Surface Plot plugin.

### Mathematical model and simulations

The minimal energy shapes of the budding tubes were determined with the finite element code *Surface Evolver*, an open-source software. Details are presented in the SI.

## Data availability

The data that support the findings of this study are available from the corresponding author upon reasonable request.

## Acknowledgements

We thank Prof. Aldo Jesorka from the Chalmers University of Technology, Sweden for his invaluable comments on the manuscript; as well as Prof. Paul G. Dommersnes from the Norwegian University of Science and Technology; and Dr. Tak Shing Chan from the University of Oslo for stimulating discussions. This work was made possible through financial support obtained from the Research Council of Norway (Forskningsrådet) Project Grant 274433, UiO: Life Sciences Convergence Environment, the Swedish Research Council (Vetenskapsrådet) Project Grant 2015-04561 as well as the start up funding provided by the Centre for Molecular Medicine Norway & Faculty of Mathematics and Natural Sciences at the University of Oslo. S.L. and A.C. gratefully acknowledge funding from the Research Council of Norway Project Grant 263056.

## Author contributions

I.G. designed research, E.K., I.K., R.O., S.L., A.C. performed research, E.K. analyzed data and E.K., S.L., A.C. and I.G. wrote the paper.

## Competing interests

The authors declare no competing financial interests.

## Additional information

## Electronic supplementary material

PDF Document including additional figures and analyses; Movies S1-7

